# Impaired dynamics of brain precapillary sphincters and pericytes at first order capillaries explains reduced neurovascular functions in aging

**DOI:** 10.1101/2021.08.05.455300

**Authors:** Changsi Cai, Stefan Andreas Zambach, Søren Grubb, Kirsten Joan Thomsen, Barbara Lykke Lind, Bjørn Olav Hald, Micael Lønstrup, Reena Murmu Nielsen, Martin Johannes Lauritzen

## Abstract

The microvascular inflow tract (MIT), comprising the penetrating arterioles, precapillary sphincters, and first order capillaries, is the bottleneck for brain blood flow and energy supply. However, the exact structural and functional alterations of the MIT during aging remain elusive. *In vivo* 4-dimensional two-photon imaging showed an age-dependent decrease in vaso-responsivity, with reduced sensitivity of the MIT to pinacidil and papaverine, vasoconstrictor endothelin-1, and nitric oxide synthase inhibitor L-NAME. This was accompanied by an age-dependent decrease in capillary density close to the arterioles and loss of pericyte processes, though the number of pericyte somas and pericyte αSMA density were preserved. The age-related reduction in vascular reactivity was most pronounced at precapillary sphincters, highlighting their crucial role in capillary blood flow regulation. Mathematical modeling revealed dysregulated but preserved pressure and flow in aged mice during vasoconstriction. Preventing reduced responsivity of the MIT may ameliorate the blood flow decrease associated with aging-related brain frailty.

---

In the aging brain, the global level of cerebral blood flow (CBF) is reduced, though the energy consumption is relatively preserved. This generates a mismatch between metabolic needs and blood supply (1, 2). In addition, neurovascular coupling (NVC), the increase in CBF that accompanies increased brain activity, is impaired, whereas the activity-dependent increases in oxygen consumption are preserved or increased (3, 4). The alterations in cerebrovascular function challenge how well the aged brain can cope with, for example, fever and excessive metabolic load associated with trauma, ischemia, or neurodegenerative processes (5, 6). Recent data suggest that pericytes fail to control key neurovascular functions in aged mice and that neurovascular dysfunction precedes age-related impairments in information processing and cognitive decline (7). However, the exact vascular mechanisms are not well understood. The objective of the present study was to provide a comprehensive dataset of the microvascular structure and function of the aged brain. This was achieved by 4-dimensional (4D; x-y-z-t) two-photon imaging of the mouse brain with a focus on the contribution of functional compartments of the microvascular inflow tract (MIT), which comprises arterioles, precapillary sphincters, and arteriolar capillaries.

NVC ensures that the local cerebral energy supply is matched to local cerebral activity and is based on a complex set of interactions between nerve cells and vascular cells. The stimulus-induced increases in local hemodynamic signals rapidly and flexibly re-allocate blood glucose and O_2_ to active nerve cells. NVC is evoked by signaling processes that, to a large extent, rely on increases in cytosolic Ca^2+^ in neurons and astrocytes (8–13), though Ca^2+^-independent processes also contribute (14–16). Brain penetrating arterioles (PAs), precapillary sphincters, and capillaries close to the arteriole are thought to play a key role. Pericytes embrace capillaries, similar to vascular smooth muscle cells, which encircle arteries and arterioles, but the pericytes close to PAs are contractile and regulate capillary blood flow (17, 18). In particular, precapillary sphincters at the junction of the PA and its initial branch (1^st^ order capillary) are crucial in maintaining capillary perfusion and are a bottleneck for brain energy supply (19), but little is known about the changes that occur in the structure and function of the MIT or this vascular segment during aging.

Here, we tested the hypothesis that impaired NVC in the aging brain is linked to alterations in vascular reactivity and pericyte coverage at specific zones of the MIT. To systematically test this idea, we studied the responsivity of the vascular segments to a suite of vasoactive compounds with known vascular effects and investigated the morphology of the MIT in aged mice using 4D two-photon microscopy in the living brain. Finally, we applied a mathematical model to provide a simple framework within which we can explain how our findings may translate to local blood flow and pressure in the microvascular network.

## RESULTS

### Capillary NVC is reduced with aging but neural activity preserved

We used NG2-DsRed mice expressing red fluorescent proteins in vascular mural cells (pericytes and vascular smooth muscle cells) under the NG2 promoter. FITC-dextran was injected intravenously to stain the vessel lumen (**Fig. 1a, b**). Using *in vivo* two-photon microscopy, we classified capillaries by their branching orders, with the 1^st^ order being the first capillary branching from the PA, and so forth (**Fig. 1b**).

**Figure 1.**
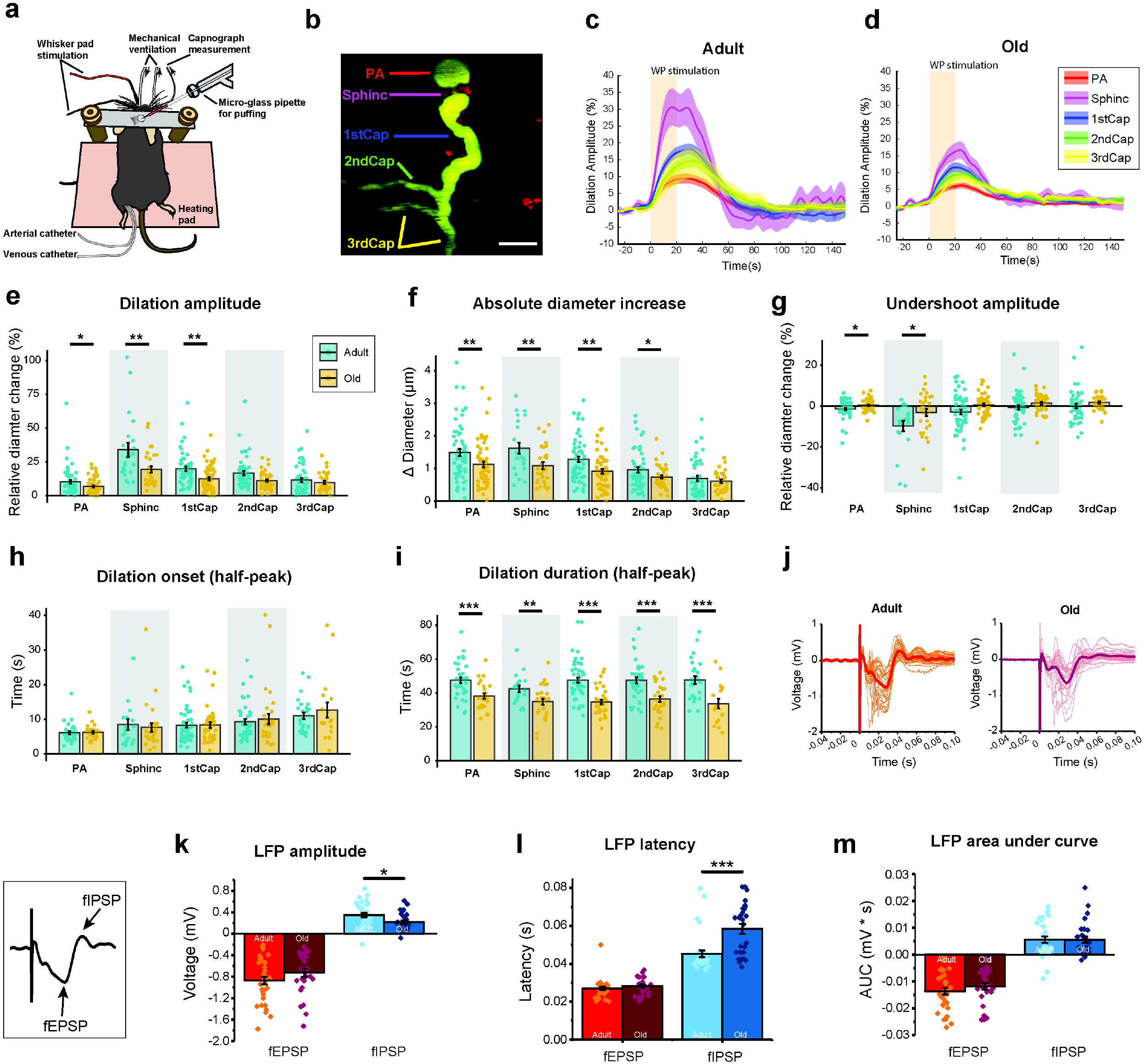
Whisker pad (WP) stimulation-induced vasodilation is reduced in aging despite preserved neural activity. (**a**) Diagram of our *in vivo* experimental setup. The physiological state of the mouse was monitored throughout the experiment, including arterial blood pressure, exhaled CO2, body temperature, heart rate, and O_2_ saturation. Two stimulations were used: WP stimulation and a glass micropipette inserted into the target area for local ejection (puff). (**b**) A two-photon image obtained by maximum intensity projection of a local image stack containing the penetrating arteriole (PA), precapillary sphincter (Sphinc), and 1^st^ – 3^rd^ order capillaries (1stCap, 2ndCap, 3rdCap). Scale bar: 10 μm. (**c**) Mean (solid curve) and SEM (shadow) traces of the vessel diameter change at each vessel location upon WP stimulation in adult and (**d**) old mice. Adult: N=31 animals, n=56 vessels. Old: N=26 animals, n=53 vessels. (**e**) Comparison of relative dilation amplitude, (**f**) absolute diameter increase by dilation, (**g**) undershoot amplitude, (**h**) half-peak dilation onset and (**i**) half-peak dilation duration in adult and old brains. (**j**-**m**) Elicited local field potentials (LFPs) by WP stimulation and recorded by the same glass micropipette. (**j**) Mean and raw LFPs in adult and old mice. (**k**) Comparison of the LFP amplitudes of both the field excitatory post-synaptic potential (fEPSP) and field inhibitory post-synaptic potential (fIPSP). (**l**) Comparison of the LFP latencies of both fEPSP and fIPSP. (**m**) Comparison of the area under the curve (AUC) of fEPSP and fIPSP in adult and old mice. Adult: N=17 animals, n=34 vessels. Old: N=17 animals, n=25 vessels. Linear mixed effect models were used to test for differences among vessel segments, followed by Tukey post hoc tests for pairwise comparisons. Data are given as mean ± SEM. * indicates p<0.05, ** indicates p<0.001, *** indicates p<0.0001.

We employed fast and repetitive 4D two-photon microscopy to examine NVC in the somatosensory cortex of anesthetized mice, minimizing the focus drift of *in vivo* imaging. Precapillary sphincters were identified as the indentation between the PA and the 1^st^ order capillary with thick mural cells enwrapping the lumen (**Fig. 1b**) (20). We used whisker pad (WP) stimulation for NVC studies. WP stimulation with 1.5 mA at 2 Hz for 20 s induced vasodilation, followed by a small undershoot at the PA, precapillary sphincter, and 1^st^ order capillary (**Fig. 1c, d**). Dilation amplitude (relative change in diameter) and absolute increases in diameter significantly declined with age (**Fig. 1e, f**). We also noted a decrease in the extent of the undershoot with age at PAs and precapillary sphincters (**Fig. 1g**). The onset of dilation (half the peak amplitude) showed no difference in adult and aged mice (**Fig. 1h**), but the dilation duration was significantly shorter at all vascular locations in old mice (**Fig. 1i**). To examine whether the reduced vascular responsivity was associated with decreased neural activity in the aged brain, we inserted a glass micropipette to record local field potentials (LFPs) (**Fig. 1a, j**). WP stimulation induced field excitatory post-synaptic potentials (fEPSPs) and inhibitory post-synaptic potentials (fIPSPs). The amplitude and latency of the fEPSP were not altered with age. In comparison, the amplitude of the fIPSP decreased and the latency of the fIPSP became prolonged in aged mice (**Fig. 1k, l**), which may suggest a failure in the synchronization of inhibitory synaptic responses in aged brains. However, the areas under the curve for both the fEPSP and fIPSP responses were similar in adult and aged mice (**Fig. 1m**), indicating preserved excitatory and inhibitory synaptic responses during aging. Overall, the mismatch between intact LFPs and the reduction in the NVC response in aged mice suggests a decrease in vascular responsivity to the preserved excitatory neurotransmission, which is the main driving force for NVC (1, 21).

### Reduced NVC in the aging brain is pericyte-dependent

The next question we addressed was whether the decrease in NVC responses with age represent a general decrease in vascular responsivity. To selectively activate pericytes, we locally puffed pinacidil, a K_ATP_ channel opener, via a glass micropipette in proximity to the imaged vessels (**Fig. 2a**). Our previous work suggested that 5 mM pinacidil acts directly on vascular mural cells without elicitation of neuronal or astrocytic responses (20). Pinacidil at 5 mM evoked vasodilation at all MIT locations: the PA, precapillary sphincter, and 1^st^ – 3^rd^ order capillaries (**Fig. 2a-c**). Compared to adult brains, PAs, precapillary sphincters, and 1^st^ order capillaries exhibited smaller dilation amplitudes and absolute diameter changes with age, whereas the responsivity was the same for 2^nd^ and 3^rd^ order capillaries (**Fig. 2d, e**). The reduced vasodilator responses in aged brains may be explained, in part, by vessel stiffness due to a reduction in the number of elastin fibers and the fragmentation of elastin, along with the deposition of collagens (22, 23). Alternatively, reduced vasodilation could be attributed to an age-related increase in vascular tone. To test this hypothesis, we puffed 10 mM papaverine on the MIT in young adult and aged mice. Papaverine dilates blood vessels by blocking vascular phosphodiesterases (24), preventing degradation of cGMP and cAMP (25). Papaverine elicited vasodilation in adult mice, but its effect in aged mice was significantly attenuated (**Fig. 2f-h**). In particular, the response to papaverine was reduced at precapillary sphincters, providing evidence of age-related changes specifically at this vascular zone.

**Figure 2.**
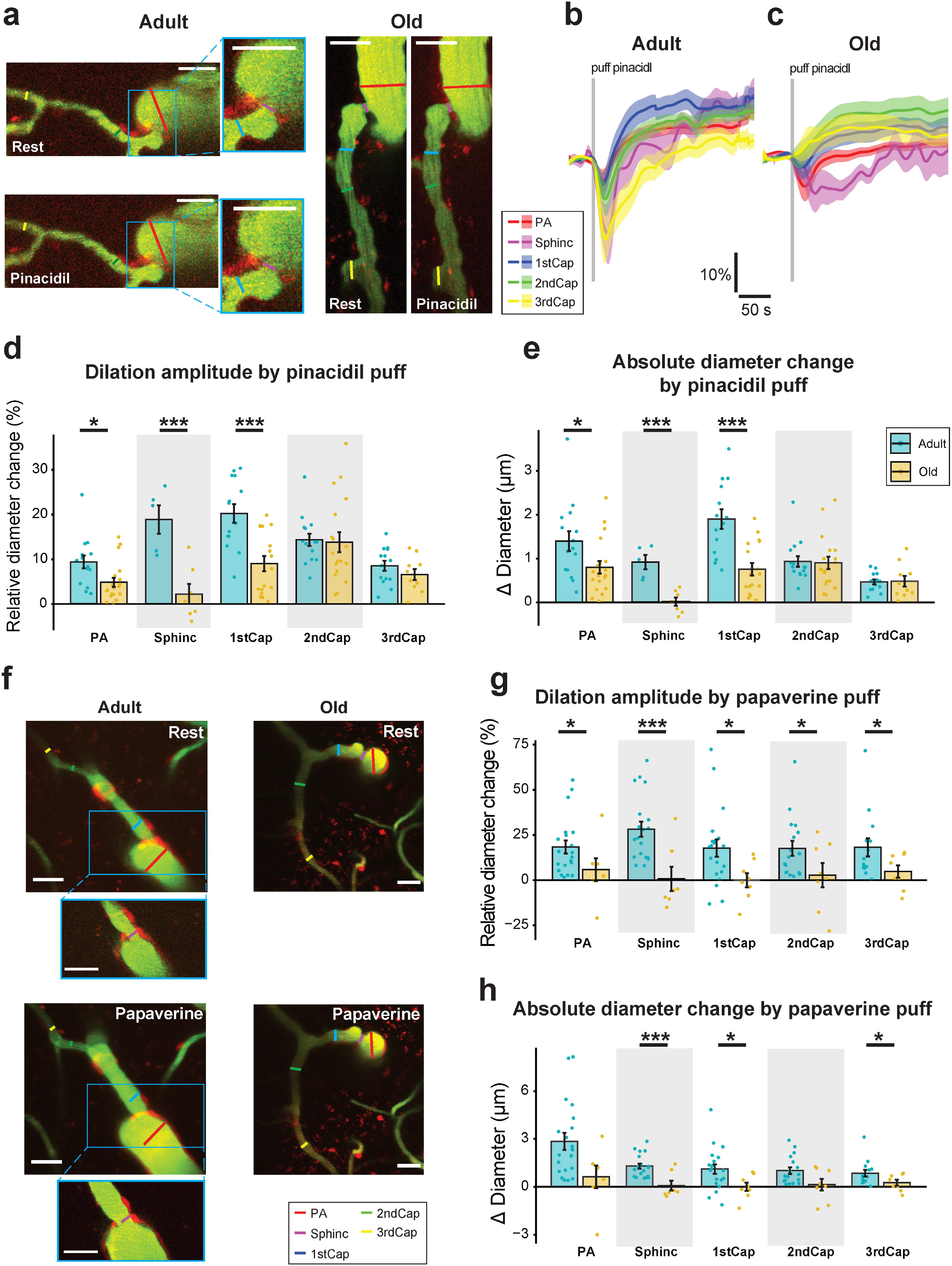
Pinacidil and papaverine-induced vasodilation is attenuated in brain aging and pericyte-dependent. (**a**) Representative images of MIT in response to 5 mM pinacidil puffing in adult and old mice. Pinacidil is an ATP-sensitive potassium (K_atp_) channel opener. Small insets show enlarged precapillary sphincters. Scale bar: 20 μm. (**b**) Mean (solid curve) and SEM (shadow) traces of the vessel diameter change at each capillary location upon pinacidil puffing in adult and (**c**) old mice. (**d**) Comparison of relative dilation amplitude and (**e**) absolute diameter change by pinacidil puffing. Adult: N=9 animals, n=15 vessels. Old: N=11 animals, n=17 vessels. Raw data for pinacidil puffing in adult brains were reused from our previous publication (20). (**f-h**) Comparison of vessel dilation with 10 mM papaverine puffing. Papaverine is a potent vasodilator that blocks vascular phosphodiesterases. Small insets show enlarged precapillary sphincters. (**f**) Representative images in response to papaverine puffing. (**g**) Comparison of relative dilation amplitude and (h) absolute diameter change with papaverine puffing. Adult: N=8 animals, n=17 vessels. Old: N=5 animals, n=8 vessels. Linear mixed effect models were used to test for differences among vessel segments, followed by Tukey post hoc tests for pairwise comparisons. Data are given as mean ± SEM. * indicates p<0.05, ** indicates p<0.001, *** indicates p<0.0001.

The vasodilation induced by local papaverine application was smaller than that induced by WP stimulation and pinacidil puffing in aged mice, but these vasodilator responses were similar in adult mice (**Supplementary Fig. 1**). As papaverine inhibits cGMP degradation, a decreased papaverine response may suggest reduced cGMP synthesis due to a reduction in nitric oxide synthesis (NOS) with age. Endothelial NOS (eNOS) activity has been reported to decline with age (26), partially due to an increase in the endogenous inhibitor of eNOS (27). To investigate whether eNOS-associated responses were altered at the MIT with age, we intravenously infused the NOS inhibitor NG-nitro-L-arginine methyl ester (L-NAME) at a dosage of 30 mg/kg. L-NAME induced vasoconstriction at the PA, precapillary sphincter, and 1^st^ – 3^rd^ order capillaries in adult mice, but to a lesser extent in aged mice (**Fig. 3a-c**). PAs and precapillary sphincters constricted less at 30 s and 150 s after L-NAME infusion in aged mice (**Fig. 3d, e**). This may corroborate the hypothesis that NO-dependent mechanisms are impaired with age.

**Figure 3.**
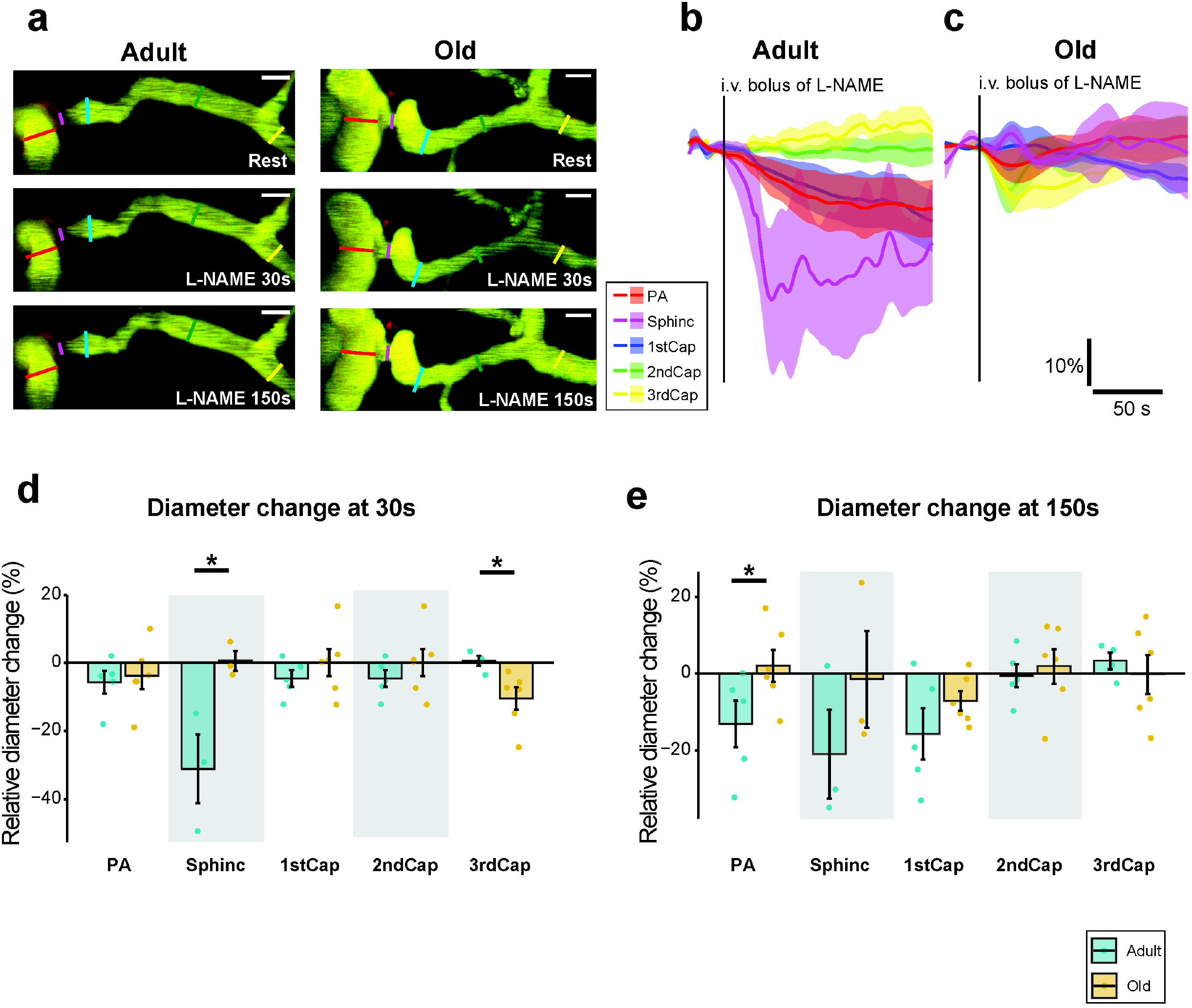
L-NAME induced vasoconstriction is decreased in penetrating arterioles and precapillary sphincters with age. (**a**) Representative images of MIT in response to intravenous bolus infusion of L-NAME at a dosage of 30 mg/kg in adult and old mice. Scale bar: 10 μm. (**b**) Mean (solid curve) and SEM (shadow) traces of the vessel diameter changes at each vascular location upon L-NAME infusion in adult and (**c**) old mice. (**d**) Comparison of diameter changes at 30 s and (**e**) 150 s by L-NAME infusion in adult and old mice. Adult: N=5 animals, n=5 vessels. Old: N=6 animals, n=6 vessels. Raw data for the intravenous infusion of L-NAME in adult brains were reused from our previous publication (20). Linear mixed effect models were used to test for differences among vessel segments, followed by Tukey post hoc tests for pairwise comparisons. Data are given as mean ± SEM. * indicates p<0.05, ** indicates p<0.001, *** indicates p<0.0001.

Taken together, the results indicate that age affects the MIT by impairing vasodilator sensitivity in the whole MIT, impairing NO-dependent mechanisms, and affecting responsivity, particularly at precapillary sphincters.

### Vasoconstriction at precapillary sphincters decreases with age

Having established that the responsivity of vascular mural cells at the MIT decrease with age, with less pronounced vasoconstriction by L-NAME infusion in aged mice, we examined whether aging affects capillary constriction. For this purpose, we used endothelin (ET1), a potent vasoconstrictor that is synthesized mainly by endothelial cells in brain pathology (28). Local puffing of 0.5 μM ET1 using glass micropipettes elicited strong and long-lasting vasoconstriction at all examined MIT locations in both adult and old mice (**Fig. 4a-c**). The contractibility of mural cells was largely preserved with age, except for precapillary sphincters, which presented reduced responsivity (**Fig. 4d, e**).

**Figure 4.**
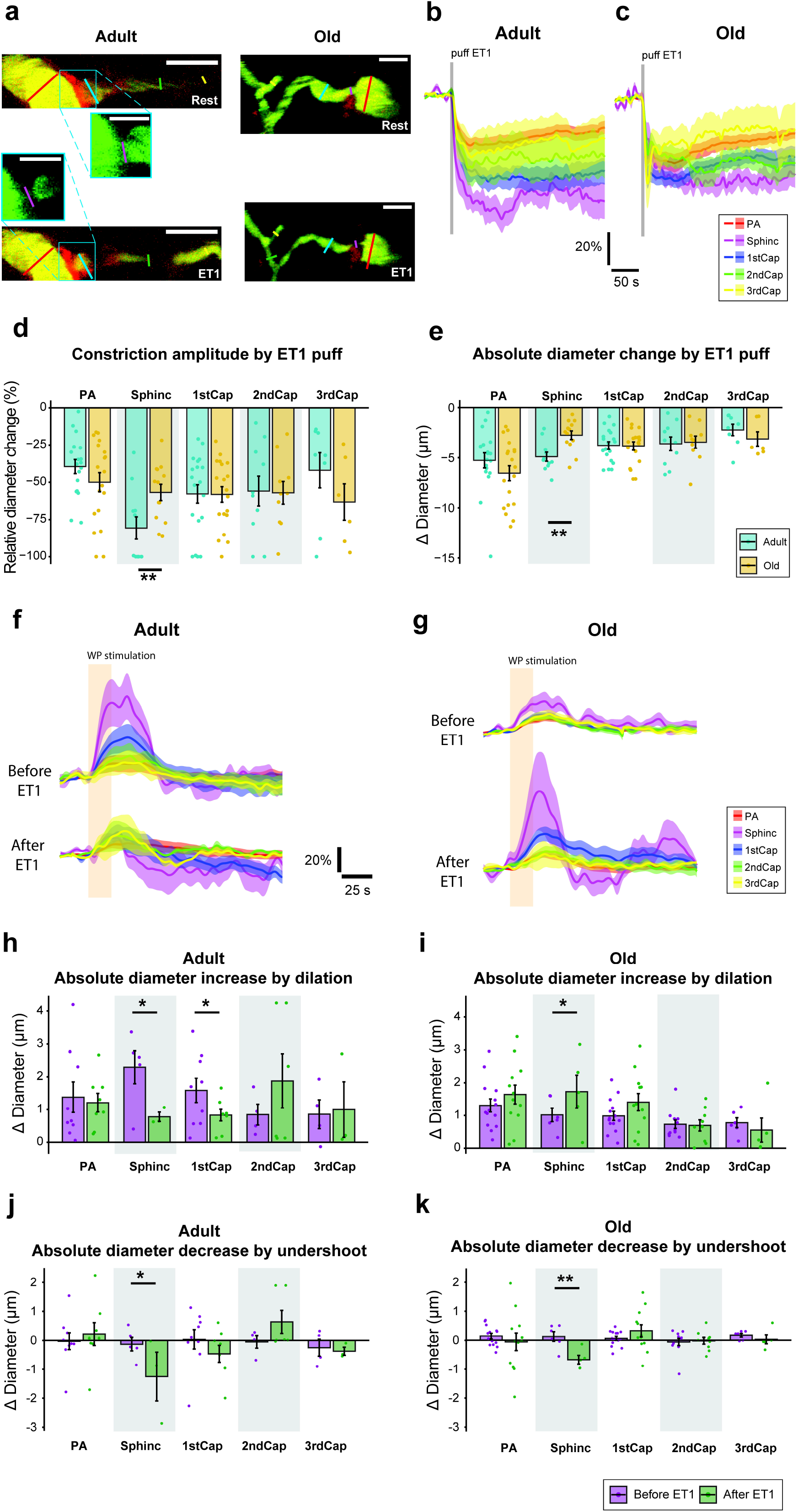
Endothelin-1 (ET1)-induced vasoconstriction decreased in pre-capillary sphincters with age. (**a**) Representative images of MIT in response to 0.5 μM ET1 puffing in adult and old mice. Scale bar: 20 μm. Small insets show enlarged precapillary sphincters at the same imaging plane in the green channel. Scale bar: 10 μm. (**b**) Mean (solid curve) and SEM (shadow) traces of the vessel diameter change at each capillary location upon ET1 puffing in adult and (**c**) old mice. (**d**) Comparison of relative dilation amplitude and (**e**) absolute diameter change after ET1 puffing. Adult: N=5 animals, n=10 vessels. Old: N=7 animals, n=15 vessels. Raw data for ET1 puffing in adult brains were reused from our previous publication (20). (**f**-**k**) Comparison of whisker pad (WP) stimulation-induced vascular responses before and after ET1 puff in adult and aged mice. (**f**) Mean (solid curve) and SEM (shadow) traces of the vessel diameter changes at each vessel location upon WP stimulation in adult and (**g**) old mice. (**h**) Comparison of the absolute diameter increase by dilation before and after ET1 puff in adult mice and (**i**) old mice. (**j**) Comparison of the absolute diameter decrease by undershoot before and after ET1 puff in adult mice and (**k**) old mice. Adult: N=5 animals, n=9 vessels. Old: N=7 animals, n=15 vessels. Linear mixed effect models were used to test for differences among vessel segments, followed by Tukey post hoc tests for pairwise comparisons. Data are given as mean ± SEM. * indicates p<0.05, ** indicates p<0.001, *** indicates p<0.0001.

Next, we asked whether NVC is altered with age because of an increase in vascular tone. For this purpose, we used ET1 again because this compound elicits a potent, long-lasting vasoconstriction, elevating local vascular tone. We recorded changes in the vascular diameter at the MIT in response to WP stimulation before and 15 minutes after 0.5 μM ET1 in both adult and aged mice (**Fig. 4f, g**). In adult mice, precapillary sphincters and 1^st^ order capillaries exhibited reduced WP-induced vasodilation after ET1 puffing. In contrast, all of the MIT of old mice presented increased WP stimulation-induced vasodilation after ET1 puffing, particularly the precapillary sphincters (**Fig. 4h-i and Supplementary Fig. 2a, b**). The following undershoots had enhanced negative amplitudes in both adult and old brains after ET1, which may be explained by increased vascular tone (**Fig. 4j-k** and **Supplementary Fig. 2c, d**). However, an increase in the vascular tone does not explain the decreased NVC in aged mice.

Therefore, with age, the PA and 1^st^ – 3^rd^ order capillaries preserve their ability to constrict, and precapillary sphincters are less contractile and dilate more at an elevated vascular tone in response to a synaptic input. This large vasodilation may serve as a compensatory mechanism for the global age-related decrease in vascular responsivity and help maintain a match between energy demand and supply in the aged brain.

### Density of pericyte processes decreases with age, but the number of pericytes and αSMA density are preserved

Capillary pericytes have been reported to degenerate during aging (29), but more detailed knowledge is lacking. To further characterize morphological changes in pericytes, we used immunohistochemistry to stain α-smooth muscle actin (αSV1A) in vascular mural cells of the MIT (**Fig. 5a, b**). The brains were prepared in the same batch to minimize variability and to ensure the comparability of antibody staining in all brain slices. For image analysis, we normalized the background of each x-y plane in one acquired image z-stack to minimize the difference in cross-plane fluorescence intensity (see details in Methods and Materials and **Supplementary Fig. 3a**).

**Figure 5.**
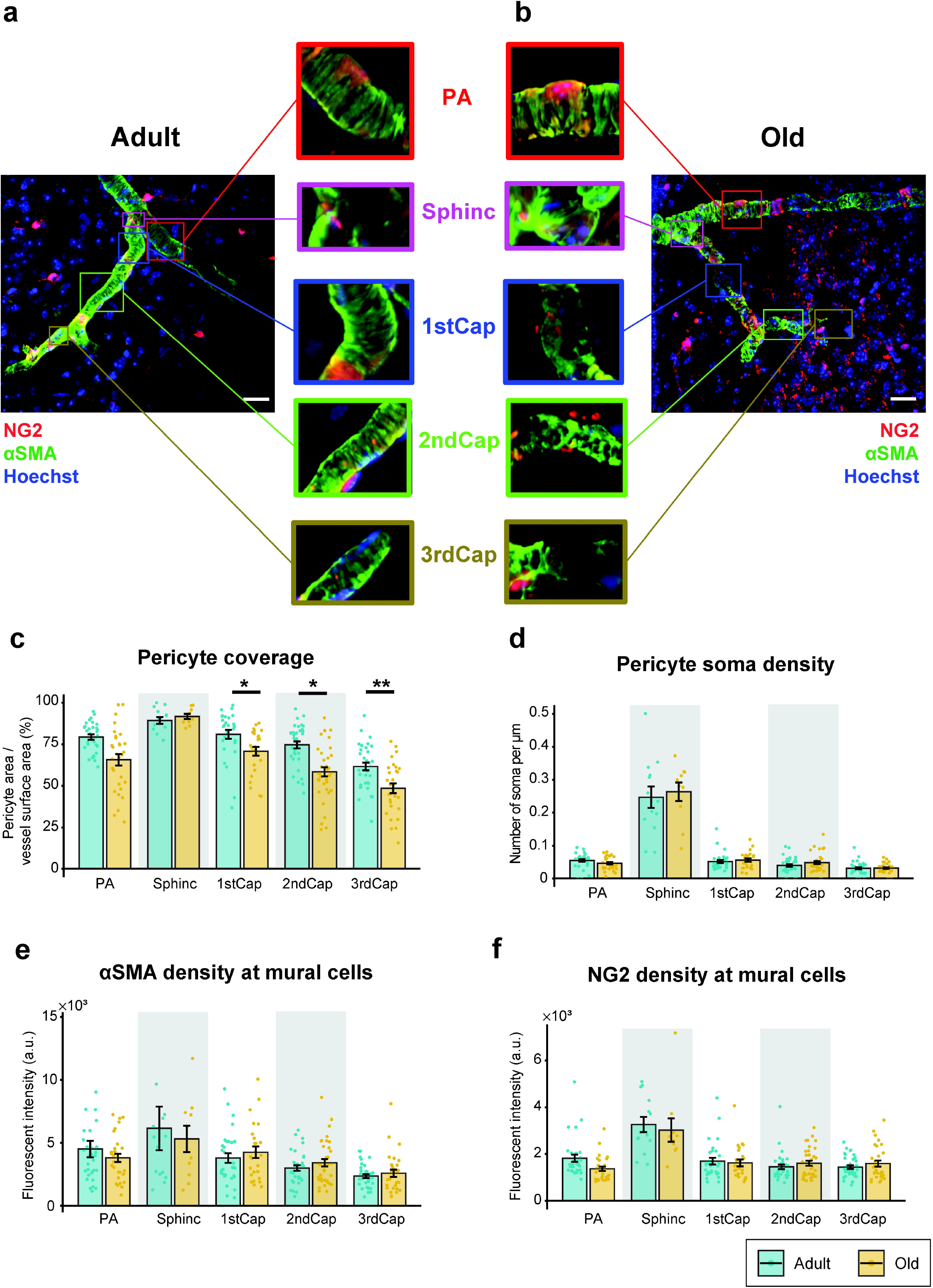
Examination of pericyte morphology and α-smooth muscle actin (αSMA) density by immunohistochemistry in adult and aged mice. (**a**) Maximum intensity projection of an adult and (**b**) old brain image stack. Red: NG2; Green: αSMA; Blue: Hoechst. Enlarged insets of each capillary segment are presented in the middle for comparison. (**c**) Pericyte coverage of vessel surface obtained by dividing pericyte area and total vessel surface area. (**d**) Pericyte soma density obtained by dividing the number of pericyte somas and the examined vessel length. (**e**) αSMA density at mural cells obtained by calculating the mean αSMA fluorescence intensity at NG2-positive vessel areas. (**f**) NG2 density at mural cells obtained by calculating the mean NG2 fluorescence intensity at NG2-positive vessel areas. Adult: N=3 animals, n=25 vessels. Old: N=3 animals, n=26 vessels. Linear mixed effect models were used to test for differences among vessel segments, followed by Tukey post hoc tests for pairwise comparisons. Data are given as mean ± SEM. * indicates p<0.05, ** indicates p<0.001, *** indicates p<0.0001.

We examined whether aging caused an alteration in pericyte αSMA or pericyte morphology at PAs, precapillary sphincters, and 1^st^ – 3^rd^ order capillaries. Pericyte coverage of the capillary surface area significantly declined at 1^st^ – 3^rd^ capillaries but was preserved at PAs and the precapillary sphincters (**Fig. 5c**). On the other hand, pericyte soma density, i.e., number of pericyte soma per millimeter of vessel length, remained constant with age (**Fig. 5d**). We also measured NG2 density and αSMA density at both mural cells (i.e., NG2-positive area) and the whole vessel surface (**Supplementary Fig. 3b**). Both NG2 and αSMA density were not altered with age (**Fig. 5e, f and Supplementary Fig. 3d, e**), suggesting that brain vascular aging is unrelated to capillary pericyte loss or αSMA degeneration, but is accompanied by lower coverage of pericyte processes.

### Capillary density declines and diameter increases with age, dependent on capillary order and cortical depth

To further characterize the morphological and structural changes of the MIT with age, we recorded two-photon image stacks consisting of PAs and their associated capillary networks in both adult and aged mice (**Fig. 6a**). Image stacks were skeletonized to estimate vessel length and diameter. We manually labeled individual PAs and 1^st^ – 5^th^ order capillaries by visual inspection (**Fig. 6b, c**). Similar to recent studies (30–32), we used total length per brain volume as a measurement of capillary density (**Fig. 6d-f**). The total length of the PA and 1^st^ – 4^th^ order capillaries significantly declined with age, whereas the total length of ≥ 5 ^th^ order capillaries was preserved (**Fig. 6e**). This implies that the capillary density declines with age close to the PA, but not on the venous side, i.e., the capillary density in the cortex is capillary-order-dependent. We also used total vessel number to estimate the capillary density and found that the total vessel number decreased with age, especially close to the PA (**Supplementary Fig. 4a-c**). Finally, we asked whether this dependency changed with depth. Our data show that the age-related decline in capillary density occurred at all cortical layers (**Fig. 6f and Supplementary Fig. 4a, b**).

**Figure 6.**
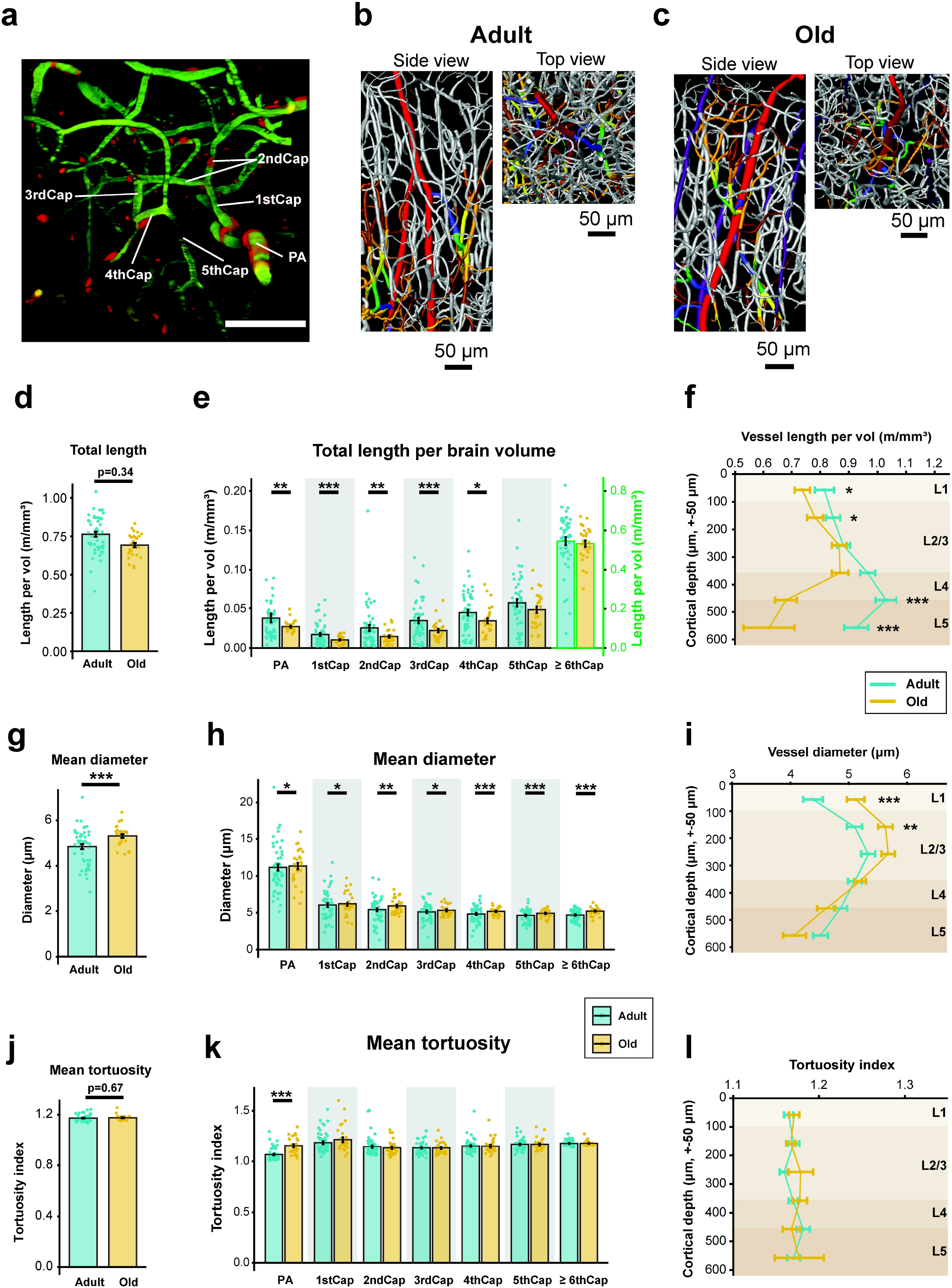
Aging induced angioarchitectural remodeling. (**a**) A representative image stack recorded by two-photon microscopy, containing the penetrating arteriole (PA), 1^st^ order capillary (1stCap), 2^nd^ order capillary (2ndCap), 3^rd^ order capillary (3rdCap), 4^th^ order capillary (4thCap), and 5^th^ order capillary (5thCap). Scale bar: 50 μm. (**b**) 3D reconstructed vasculature of the somatosensory cortex in an adult mouse and (**c**) old mouse. Raw data were obtained by two-photon microscopy and analyzed by Amira software. Left: Side view. Right: Top view. Red: PAs; Blue: 1stCap; Green: 2ndCap; Yellow: 3rdCap; Orange: 4thCap; Brown: 5thCap; Purple: ascending venules. (**d**-**f**) Vessel total length per brain volume in adult and old somatosensory cortex. (**d**) General vessel total length, (**e**) capillary order dependent vessel total length, and (**f**) cortical depth dependent vessel total length. (**g**-**i**) Vessel mean diameter in adult and old somatosensory cortex. (**g**) General mean diameter, (**h**) capillary order dependent mean diameter, and (**i**) cortical depth dependent mean diameter. (**j**-**l**) Vessel mean tortuosity in adult and old somatosensory cortex. The tortuosity index denotes division of the curved length by the chord length of each vessel segment. (**j**) General mean tortuosity, (**k**) capillary order dependent mean tortuosity, and (**l**) cortical depth dependent mean tortuosity. Adult: N=6 animals, n=52 vessels. Old: N=4 animals, n=29 vessels. Linear mixed effect models were used to test for differences among vessel segments, followed by Tukey post hoc tests for pairwise comparisons. Data are given as mean ± SEM. * indicates p<0.05, ** indicates p<0.001, *** indicates p<0.0001. L1 – L5 denote cortical sublayers.

Consistent with previous reports (2, 33, 34), vessel diameter increased with age for all capillary orders (**Fig. 6g, h**), but this was only significant at a depth of 0 – 200 μm (**Fig. 6i**). Considering that the mean vessel diameter increased and total vessel length decreased with age, we examined the total vessel volume (**Supplementary Fig. 4d-f**) as a proxy for brain perfusion capacity. Interestingly, the global vessel volume increased with age (**Supplementary Fig. 4d**) as a result of an increase in the volume of the venous capillaries (**Supplementary Fig. 4e**), which suggests that brain vascular aging is associated with an increase in venous compliance. Lastly, we estimated vessel tortuosity in both adult and old brains by dividing the curved length by the chord length of each vessel segment (tortuosity index) (**Fig. 6j-l**). The PAs became significantly more tortuous with age, whereas vessels downstream from the PA remained at the same level of tortuosity (**Fig. 6k-l**).

Taken together, the results suggest that capillary density decreases close to the PA, capillary total volume increases close to cortical venules, the capillary changes are similar for cortical layers I-III, and vessel tortuosity increases at the PA, but not in capillaries.

### Capillary blood flow and pressure are affected by aging

To assess the possible implications of a special role of the sphincter and 1^st^ order capillary on capillary flow and pressure, we developed a mathematical model based on 3-dimensional reconstruction of the brain vasculatures from three adult and three old mice, including all elements of the MIT and up to the 6^th^ order capillary (**Fig. 7a, b**). The model assumed that the vessels were rigid and the flow laminar. We calculated the flow resistance at each capillary order using Poiseuille’s law. Given that the sum of flows entering and leaving any internal node equals zero (Kirchoff s 2^nd^ law), we calculated the blood pressure and flow at each node within the vascular network using Ohm’s law. We designated vessel diameters and dilation and constriction amplitudes based on our measurements by two-photon imaging. We calculated how blood flow and pressure in specific vessel segments changed from the resting state (control) to WP stimulation (vasodilation) and ET1 puffing (vasoconstriction).

**Figure 7.**
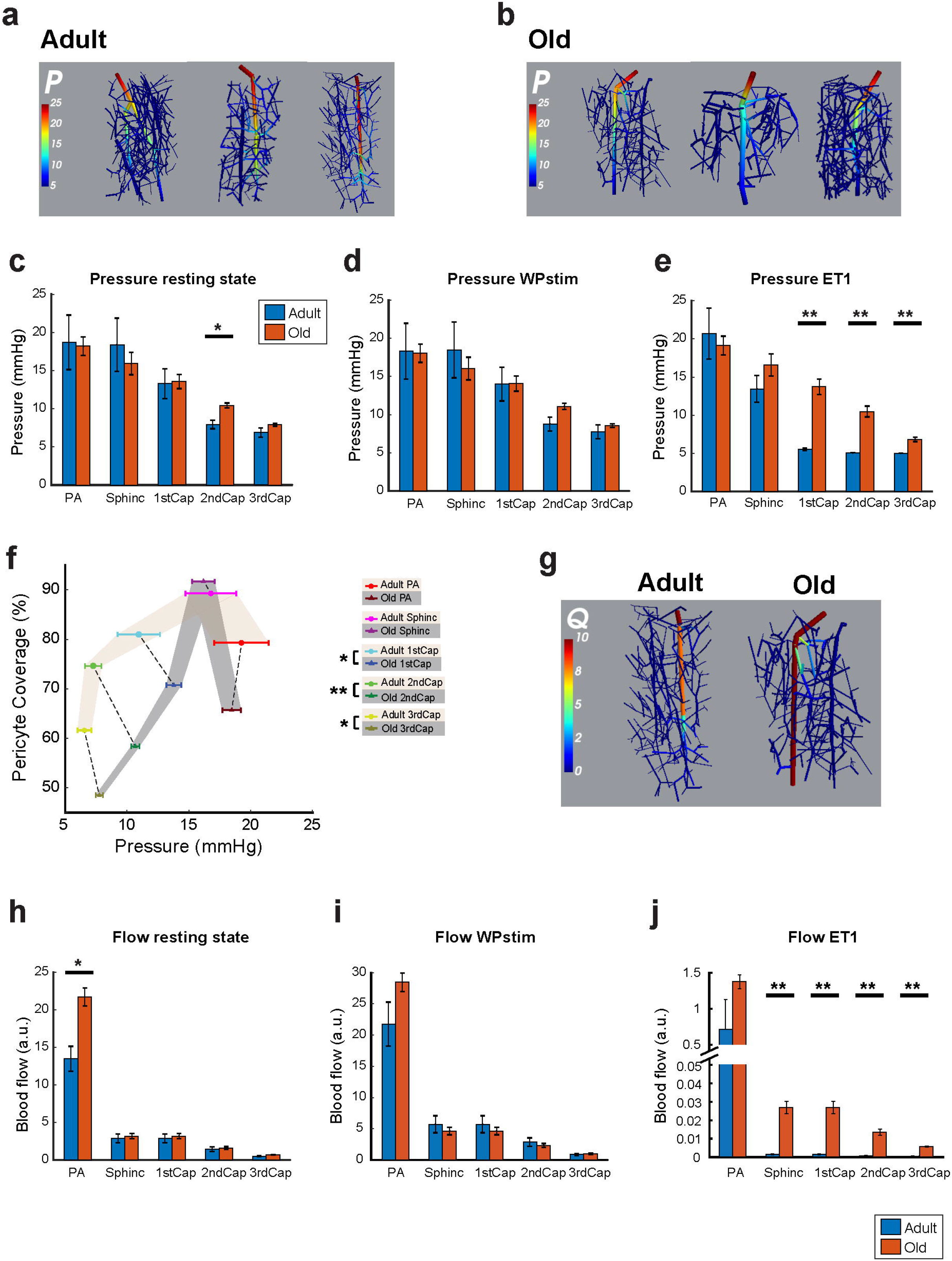
Math modeling indicates that vascular blood flow and pressure are affected by aging. (**a**) Mathematical modeling of pressure distribution in 3D reconstructed vasculature from 3 adult and (**b**) 3 old mouse brains. (**c**) Summary of the average pressures in penetrating arteriole (PAs), precapillary sphincters, and 1^st^ – 3^rd^ order capillaries in the networks under resting-state, (**d**) with whisker pad (WP) stimulation, and (**e**) after ET1 puff. (**f**) Correlation of pericyte coverage and blood pressure summarized for all three states. The dashed line connects the adult and old measurements at the same vascular location. (**g**) Mathematical modeling of flow distribution in 3D reconstructed vasculature from 3 adult and 3 old mouse brains. Only one representative image from each group is shown. (**h**) Summary of the average flow in PAs, precapillary sphincters, and 1^st^ – 3^rd^ order capillaries in the networks under resting-state, (**i**) WP stim, and (**j**) after ET1 puff. Adult: N=3 animals, n=3 vessels; Old: N=3 animals, n=3 vessels. Linear mixed effect models were used to test for differences among vessel segments, followed by Tukey post hoc tests for pairwise comparisons. Data are given as mean ± SEM. * indicates p<0.05, ** indicates p<0.001, *** indicates p<0.0001.

We calculated the pressure distribution in the adult and aged vascular network models in the resting state (**Fig. 7c**) and during WP stimulation (**Fig. 7d**) and ET1 puffing (**Fig. 7e**). Our model suggested that, in old mice, 1^st^ – 3^rd^ order capillaries (but not the PA or sphincter) were exposed to higher intraluminal pressure than in adult mice, particularly when exposed to ET1 (**Fig. 7c-e**). To examine whether pressures at all three states (rest, WP, and ET1) were related to the degree of pericyte coverage reported in **Fig. 5**, we plotted pericyte coverage as a function of blood pressure for adult and old mice (**Fig. 7f**). The analysis suggests that the capillary pressure distribution correlates with pericyte coverage in an almost linear manner from sphincters to the 3^rd^ order capillaries. With age, pericyte coverage decreased, whereas pressure increased at the 1^st^ – 3^rd^ order capillaries, suggesting a more fragile regulation of capillary pressure by aged pericytes (35).

Next, we compared the blood flow distribution in adult and old mice at each vessel location (**Fig. 7g-j and Supplementary Fig. 5**). Given that the mean capillary diameter was larger in the aged brain than in the adult brain, the flow at PAs was higher in old mice than in young adult mice at rest (**Fig. 7h**). Although the age-related flow difference was insignificant for WP stimulation (**Fig. 7i**), the flow drop was mitigated at precapillary sphincters and 1^st^ – 3^rd^ order capillaries in old brains after ET1 puffing (**Fig. 7j**).

Therefore, with age, the regulation of capillary pressure is more fragile. In contrast, capillary flow is maintained more stably in old mice during vasoconstriction compared to adult mice.

## DISCUSSION

Using *in vivo* 4D two-photon microscopy, immunohistochemistry, and mathematical modeling, we examined the structural and functional changes that occur in brain vessels with age. We focused on MIT, i.e., the PAs, precapillary sphincters, and first few orders of capillaries, which have been suggested to play important roles in regulating capillary blood flow. Our study shows that, at the MIT, vascular responsivity decreases with age, which contributes to the reduction in NVC responses in aged mice. In addition, the decrease in vascular dilation may be ascribed to an impaired ability of pericytes to relax, especially precapillary sphincters. In comparison, the ability of vessels to constrict is preserved with age, except for precapillary sphincters, which constrict less than in young adult mice. The reduced NVC is not explained by an increase in vascular tone because of compensatory mechanisms of vascular segments in the MIT of aged mice. Coverage of MIT by pericyte processes is reduced with age, but pericyte soma density and αSMA density remain unaltered. Capillary density decreases close to the PA, but vessel diameters increase at all locations, reducing the overall vascular resistance. Mathematical modeling suggested impaired regulation of blood pressure at capillaries but implied preserved blood flow at capillaries during vasoconstriction. Thus, our findings provide mechanistic insights into the role of precapillary sphincters and the first few orders of capillaries in brain vascular aging.

Aging is associated with a decrease in hematocrit, blood oxygenation, global CBF, capillary density, and brain tissue pO_2_, and an increase in pial artery tortuosity, capillary diameter, cerebral capillary RBC velocity, and the heterogeneity of capillary RBC transit time (1, 2, 34, 36, 37). All of these changes are accompanied by unchanged CMRO_2_ (2) and, in the aged brain, the relationship between blood flow and metabolism is complex. A preserved brain vascular supply is required to ensure the delivery of enough glucose and oxygen to support information processing and cognition. In neurodegenerative diseases, the aging-related impairment of K_ATP_ channels and NO-signaling pathways, neurovascular uncoupling, and pericyte process rarefaction and decrease in capillary density are further exacerbated (38–40). In this study, we report preserved neuronal activity in aged brains upon WP stimulation (**Fig. 1j-m**), which suggests that the vascular changes are not accompanied by changes in information processing in healthy aging.

Our 3D reconstruction of brain vascular networks revealed a decrease in capillary density close to the PA, most prominently at 1^st^ – 4^th^ order capillaries (**Fig. 6d-f and Supplementary Fig. 4a-c**). This is consistent with a previous study showing that the first few orders of capillaries have higher probability of spontaneous obstruction and subsequent pruning without compensatory sprouting by aging (41). Our findings may also be attributed to impaired arteriogenesis and angiogenesis due to a decrease in vascular endothelial growth factor (VEGF) in the aged brain (42, 43).

The decrease in total vessel length and total vessel number suggests that the maximal distance from capillary to neuron is longer in aged brains than in young adult brains. This is expected to cause longer O_2_ diffusion distances from blood vessels to brain tissue, with important implications for the brain’s energy supply. Based on our modelling work (**Fig. 7h**), we assume that the capillary blood flow at rest is not altered with age. However, the combination of circumstances could result in spatiotemporal partitioning of glycolytic and oxidative metabolism during increases in local activity, compromised delivery of oxygen, and the development of hypoxia at the sites most distant to vessels (2). This hypoxia could be exacerbated further by functional activation of the brain, which causes an increase in the neuronal activity and CMRO_2_, whereas aged brains do not receive sufficient energy supply due to the decrease in capillary dilation compared to the adult brain.

The prevalence of the precapillary sphincter is highest in the most superficial layers of the cortex (L1 – L2) (19). In aged mice, all vessel segments in the superficial layers of the cortex are increased in diameter. We speculate that the precapillary sphincter could actively protect downstream capillaries from high blood pressure increases. In this study, we observed that the function of precapillary sphincters decreases with age. This age-related decrease in precapillary sphincter function may result in less protection of downstream capillaries against high blood pressure, leading to capillary dilation. Loss of vessel tone by age-related autoregulatory dysfunction of the cerebral microvasculature may cause this increase in capillary diameter (44, 45).

In patients, isocapnic hypoxia opens K_ATP_ channels and causes an increase in the CBF, thereby maintaining oxygen delivery and brain partial O_2_ (46). With age, the increase in CBF produced by K_ATP_ channel opening is reduced, and this is further aggravated at an early stage of AD and likely caused by reduced expression of K_ATP_ channels (38). Long-term treatment of AD mice at an early stage with K_ATP_ channel opener improves cognitive function and reduces the tau and amyloid burden (47). Our results from aged mice suggest that the capillary vasodilation caused by K_ATP_ channel opener pinacidil was markedly decreased at the precapillary sphincters, 1^st^ order capillaries, and PAs, which indicates that K_ATP_ channel dysfunction contributes to impaired dysregulation of the MIT in aging, suggesting that it has clinical interest as a novel target for pharmacological treatment.

NO signaling decreases with age, which entails a reduction in the inhibition of ET secretion (48, 49), which may induce collagen formation, causing arterial stiffness and promoting atherosclerosis (50) and neurodegeneration (51). Following subarachnoid hemorrhage and stroke, ET levels are pathologically elevated and may cause vasospasms (28, 52, 53), further pointing to a pathological role of ET *in vivo*. Our results showed that vasoconstriction by ET1 was the same in young adult and aged mice, except for the precapillary sphincters. The aged sphincters constricted less than that in adult mice in response to ET1.

Immunohistochemistry of pericyte morphology and αSMA density showed preservation of αSMA density in mural cells of the MIT with age, whereas mural cell coverage by pericyte processes was decreased at 1^st^ – 3^rd^ capillaries, probably due to diminished pericyte outgrowth from aged pericytes compared to their age-matched counterparts. However, the pericyte soma density was preserved, suggesting that this is a reduction in pericyte complexity more than pericyte numbers, which is similar to neuronal changes (54). Pericyte process rarefaction could contribute to the dysregulation of blood flow at 1^st^ – 3^rd^ capillaries, as observed in our *in vivo* experiments. Furthermore, the decrease in pericyte processes may impact the blood-brain barrier (55), and pericyte loss is associated with a severe loss of blood flow and induction of the neurodegeneration cascade (56). The age-related impairment of blood flow regulation was most pronounced at precapillary sphincters, which are encircled by a mural cell, but mural cells at sphincters do not have long fine processes and the reduction in responsivity at this site is not explained by the decrease in vessel coverage by pericyte processes, but by other unknown mechanisms.

Adaptation to healthy aging may potentially be protective, but the mismatch of energy supply and demand is likely a weak point in brain vascular aging. However, the reduced NVC may limit the inflow of O_2_ to the activated brain, limiting free radical production and consequently delaying further aging (57). The decreased responsivity and increase in vascular tone with age protect the brain from the increased amplitude of the pressure pulse with every heartbeat that impacts and disturbs neuron function (58). On the other hand, the low global blood flow and reduced responsivity reduce the safety of the energy supply with age. This is a challenge in pathology when repair mechanisms are needed.

In summary, our data provide mechanistic insights into the integrated vascular functions that regulate brain capillary blood flow in the aged brain. In particular, we unveiled important roles of the precapillary sphincters and first order capillaries in healthy brain aging.

## DATA AVAILABILITY

The data that support the findings of this study are available from the corresponding author upon request.

## CODE AVAILABILITY

The custom code that support the findings of this study are available from the corresponding author upon request.

## ACKNOWLEDGEMENTS

We would like to acknowledge the Core Facility for Integrated Microscopy, Faculty of Health and Medical Sciences, University of Copenhagen, where we used spinning disc confocal microscopy in our *in vitro* studies.

This study was supported by the Lundbeck Foundation, the Danish Medical Research Council, the Alice Brenaa Foundation, Augustinus Foundation, Carl og Ellen Hertz Familielegat, A. P. Møller Foundation, The NOVO Nordisk foundation, and a Nordea Foundation Grant to the Center for Healthy Aging.

## MATERIALS AND METHODS

### Animal handling

All procedures were approved by the Danish National Ethics Committee according to the guidelines set forth in the European Council’s Convention for the Protection of Vertebrate Animals used for Experimental and Other Scientific Purposes and are in compliance with the ARRIVE guidelines. A total of 63 NG2DsRed mice Tg (cspg4-DsRed.T1)1Akik/j from Jackson laboratory, including 32 adult mice (aged 3 – 12 months) and 31 old mice (aged 20 – 28 months), were used. The mouse trachea was cannulated for mechanical ventilation. One catheter was inserted into the left femoral artery to monitor blood pressure and blood gases; a second catheter was placed into the left femoral vein for infusion of substances. To maintain the mouse under physiological conditions, we continuously monitored end-expiratory CO_2_, blood pressure, heart rate, and O_2_ saturation at the right hind paw. Throughout the experiment, we assessed blood gases in arterial blood samples twice (55 μL each time): after completion of the craniotomy and after termination of the experiments just prior to euthanasia. We monitored the mouse’s physiological state at pO_2_ 95 – 110 mmHg, pCO_2_ 30 – 40 mmHg, and pH 7.35 – 7.45. Body temperature was maintained at 37°C using a rectal temperature probe and heating blanket. We drilled a 4-mm-diameter craniotomy above the sensory barrel cortex region, centered 0.5 mm behind and 3 mm to the right of bregma. After dura removal, the preparation was covered with 0.75% agarose gel, further submerged under artificial CSF (aCSF; NaCl 120 mM, KCl 2.8 mM, NaHCO_3_ 22 mM, CaCl_2_ 1.45 mM, Na_2_HPO_4_ 1 mM, MgCl_2_ 0.876 mM, and glucose 2.55 mM; pH 7.4), and kept at 37°C. For imaging experiments, three quarters of the craniotomy was covered with a tilted glass coverslip that permitted the insertion of glass micropipettes. A detailed description was published previously (59). The mice were anesthetized by intraperitoneal injection of xylazine (10 mg/kg) followed by ketamine (60 mg/kg), and maintained during surgery with supplemental doses (30 mg/kg) of ketamine every 24 minutes. Upon completion of all surgical procedures, the anesthesia was switched to continuous intravenous infusion with a mixture of 17%α-chloralose and 2% fluorescein isothiocyanate (FITC)-dextran (0.02 mL/10 g/h). At the end of the experimental protocol, the mice were euthanized by intravenous injection of 0.05 mL pentobarbital followed by cervical dislocation.

### Whisker-pad stimulation

The mouse sensory barrel cortex was activated by stimulating the contralateral ramus infraorbitalis of the trigeminal nerve via a set of custom-made bipolar electrodes inserted percutaneously. The cathode was positioned according to the hiatus infraorbitalis (IO), and the anode was inserted into the masticatory muscles. WP stimulation (thalamocortical IO stimulation) was performed at an intensity of 1.5 mA with pulse duration of 1 ms, duration of 20 s, and frequency of 2 Hz.

### Local ejection (puffing) by glass micropipette

We used a pipette puller (P-97, Sutter Instrument) to produce borosilicate glass micropipettes with a resistance of 3 – 3.5 MΩ. The pipette was loaded with a mixture of 10 μM Alexa 594 and vasoactive substances of interest, which enabled visualization of the pipette tip under an epi-fluorescent camera and two-photon microscope. Guided first by epi-fluorescent imaging, and then by two-photon microscopy, the pipette was carefully inserted into the cortex and moved in proximity of the targeted vessels. The distance between the pipette tip and the vasculature was approximately 30 – 50 μm. Pressure ejection of vasoactive substances was achieved using an air pressure pump at 8 – 15 psi. A red cloud (Alexa 594) ejected from the pipette tip was visually observed to cover the local vascular region instantaneously, and the background returned to normal approximately 1 minute after puffing (60). An estimated volume of 0.38 nL was delivered in each puff. This high concentration is rapidly diluted. The drug concentration in the puffing solution was higher than the concentration of the same drug used in brain surface superfusion experiments. An electrode inside the puffing pipette was used to record local electrical brain activity during the experiments. This ensured that LFPs were recorded at the location where the concentration of the puffed drug was the highest.

### Two-photon imaging

FITC-dextran (MW 500,000, 50 μL, Sigma-Aldrich) was applied intravenously to stain the vessel lumen green. Before two-photon imaging, 4%w/v FITC-dextran was administered as a bolus into the femoral vein. During two-photon imaging, 2% FITC-dextran was continuously infused at a slow speed to compensate for the metabolic loss of FITC-dextran. Fast repetitive hyperstack imaging (4D imaging) was performed using a commercial two-photon microscope (Femto3D-RC, Femtonics Ltd.) and a 25 × 1.0 NA piezo motor objective. This method compensates for focus drift and allows for evaluation of the vasculature spanning a certain z-axis range. Each image stack was acquired within 1 second and comprised 10 – 14 planes with a plane distance of 4 – 5 μm. This approach covered the whole z-axis range of the investigated blood vessels. The pixel sizes in the x-y plane were 0.2 – 0.38 μm. The excitation wavelength was set to 900 nm. The emitted light was filtered to collect red and green light from DsRed (pericytes) and FITC-dextran (vessel lumens), respectively.

### Two-photon imaging data analysis

The imaging analytical tool was custom-made using MATLAB. To measure vessel diameter in the hyperstack video, we flattened each image stack onto one image by maximal intensity projection, creating a 2D time-lapse movie. An averaged image over time from the green channel was plotted for placement of regions of interest (ROIs). Rectangular ROIs with a width of 4 μm were drawn perpendicular across the vessel longitude. The rectangular ROI was averaged by projection into one line for each frame, representing the profile of the vessel segment at this frame. The profile line was plotted as a 2D image with the x axis as the number of frames. Chan-Vese segmentation or pixel intensity-based segmentation was used to delineate the vessel edge and estimate the diameter change in the vessel segment at this ROI. Less than 2% of the evoked diameter change was considered nonresponding but included in the analysis as zero values. Up to five ROIs with an adjacent distance of 10 μm were placed at the same order capillary depending on the length of the capillary order.

Diameter curves at the same order capillary were averaged, representing the response curve at this capillary order. The relative response amplitude was defined as the largest vasodilation/constriction amplitude (as a percentage) after WP stimulation/puffing. The response latency was defined as the latency of the half-max amplitude. The response duration was defined as the period between half-peaks of rising and falling phases. Stimulation and puffing were performed following 1 minute of imaging in the resting state.

The evoked LFPs were recorded and measured using the software Spike2 v7.02a (CED, Cambridge Electronic Design). LFPs were overlaid and averaged by aligning the stimulation onset time. The evoked negative field potential was identified as the fEPSP and the following positive potential as the fIPSP. The fEPSP and fIPSP amplitudes were defined as the peak amplitudes from baseline, and latency was defined as the time from stimulation onset to the peak of the fEPSP and fIPSP.

### 3D vascular reconstruction and analysis by Amira

In a dedicated set of *in vivo* experiments, we recorded z-stacks in a 250 – 500 × 250 – 500 μm area in steps of 1 – 2 μm around as many PAs in the craniotomy as possible and as deep as 600 μm below the pial surface. For this purpose, we used FITC-injected adult or aged NG2-dsRed or c57bl/6 mice. The laser wavelength was set to 920 nm, and the laser power was gradually increased with imaging depth until the signal/noise ratio was too high to resolve the vessels. We did not observe any indications of laser damage, or even bleaching of the NG2-DsRed caused by the z-stack recordings. The angioarchitecture was analyzed by skeletonization in Amira (Thermo Fisher) after applying a gaussian filter and local thresholding. We manually cleaned the skeleton from runts (small terminal branches that were obvious mistakes) and labeled the arteries and venules before incrementally labeling the 1^st^ – 5^th^ order capillaries. The capillaries were always labeled the lowest order possible. The data were exported from Amira and analyzed in Matlab using a custom-made algorithm.

### Drug application

We locally ejected vasoactive substances (pinacidil, papaverine, endothelin) via a glass micropipette. Pinacidil monohydrate (P154, Sigma Aldrich) and papaverine (P3510, Sigma Aldrich) were solubilized in dimethylsulfoxide and further dissolved with aCSF. ET1 (Cat. No. 1160, Tocris) was directly dissolved in aCSF. Aliquots were stored at −20°C. L-NAME (N5751, Sigma Aldrich) was dissolved in saline and stored at −20°C. L-NAME was intravenously injected as a bolus at a dosage of 30 mg/kg. Before each experiment, each drug solution used for puffing was mixed 1:1 with 20 μM Alexa 594 (dissolved in aCSF). Concentrations in the final puffing solution were as follows: pinacidil 5 mM, papaverine 10 mM, and ET1 0.0005 mM.

### Immunohistochemistry

Three adult and three old NG2-dsRed mice were transcardially perfused with 4% paraformaldehyde and their brains extracted and stored in paraformaldehyde for 4 hours. The brains were then cryoprotected in phosphate-buffered saline (PBS) with 30% sucrose and 0.1% sodium azide for 48 hours, rapidly frozen on dry ice-chilled isopentane, trimmed to only include the somatosensory cortex, and sectioned into coronal slices 50-μm-thick using a cryostat. Sections were rinsed in 0.1 M PBS for 5 minutes three times. The sections were permeabilized and blocked in 0.5% Triton-X 100 in 1 × PBS (pH 7.2) and 1% bovine serum albumin overnight at 4°C. The sections were further incubated for two nights at 4°C with mouse ACTA2-FITC antibodies (1:200; Sigma, F3777) in blocking buffer containing 1-5% bovine serum albumin in 0.25 – 0.5% Triton-X 100 in 1× PBS. The sections were then washed in 0.1 M PBS for 5 minutes three times and incubated in Hoechst (1:6000) for 7 minutes, rinsed again (3×5 min) in blocking buffer and two times in 1× PBS, and mounted using SlowFade™ Diamond Antifade Mountant (Invitrogen, S36963). We imaged the stained and mounted brain samples by spinning disc microscopy using a Zeiss CellObserver microscope equipped with Yokogawa spinning discs, Hamamatsu Orca Fusion scientific CMOS camera, and a Fluar 40x/1.20 oil immersion objective. For the green channel (FITC), an illumination wavelength of 488 nm and emission filter BP525/50 were used. For the red channel (DsRed), an illumination wavelength of 561 nm and emission filter BP629/62 were used. For the blue channel (Hoechst), an illumination wavelength of 405 nm and emission filter BP450/50 were used. Z-stack images of the continuous vasculature, PA, precapillary sphincters, and 1^st^ – 3^rd^ order capillaries, and if possible higher order capillaries, were recorded at a resolution of (x) 0.163 μm/pixel × (y) 0.163 μm/pixel × (z-step) 0.28 μm in coronal brain slices from the somatosensory cortex.

### Image analysis of immunohistochemistry

The existing problem with using immunohistochemistry to quantify protein density is the uneven fluorescence intensity of antibodies in brain slices from different depths due to the decline in antibody binding efficacy towards the center of the slice, as well as the laser power reduction with imaging depth. Therefore, we implemented a new image processing algorithm to overcome these problems. The same background area in each plane was manually selected from the entire image stack (**Supplementary Fig. 2a left**). Next, we calculated the mean background intensity of each plane across the entire stack, and normalized every image plane to the background peak value (**Supplementary Fig. 2a middle**). The new image stack was then flattened by maximal intensity projection (**Supplementary Fig. 2a right**). We used two different methods to quantify the αSMA intensity: the whole vessel area in the αSMA image, and the selected region in the αSMA image based on the pericyte-covered area in the DsRed image (**Supplementary Fig. 2b**).

### Computational modeling

We developed a simple hemodynamic network model based on image reconstruction of a single PA and venule and their associated capillaries. Briefly, Amira software (version 6.1) was used to segment a z-stack of images, preprocessed with a Gaussian filter and background subtraction to facilitate segmentation. Network nodes, edges, and vessel lengths and radii were extracted from the reconstruction. However, the vessel radii of the upper PA and the 1^st^, 2^nd^, and 3^rd^ order capillaries were taken from the original image data and, for each simulation, the vessel radii of specific vessels in the network were deliberately changed. Assuming that the vessels are rigid and the flow (*Q*) laminar, the flow resistance of individual vessel segments was calculated using Poiseuille’s law and the standard hemodynamics measure of flow resistance (*R*): 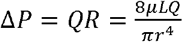, i.e., 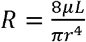, where *P* is pressure, *μ* is dynamic viscosity, *L* is vessel length, and *r* is vessel radius. In a network, Kirchoff’s law states that the sum of flows entering and leaving any internal node equals zero: 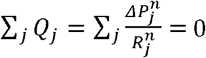, where 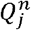 is the flow, 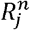 is the vascular flow resistance, and 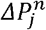 is the pressure drop in the *j*’th vessel entering the *n*’th node. We applied the empirical model describing the changes in the apparent viscosity of blood (μ) with diameter (D) and hematocrit (69, 70):

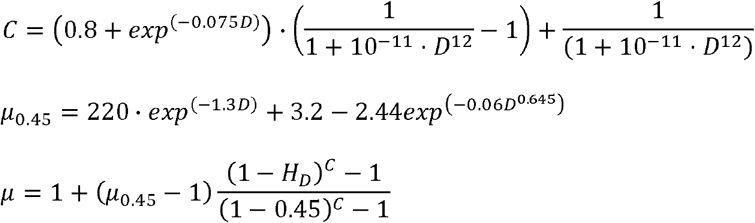

The discharge hematocrit (H_d_) was calculated based on a tube hematocrit (H_t_) of 0.3 (71):

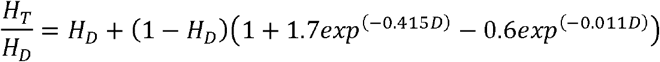

To solve the system of linear equations, we chose the boundary conditions such that the inlet pressure into the PA was 25 mmHg in the control situation and every outlet pressure of the capillaries was 5 mmHg. The system was solved using the root solver in SciPy (1.1.0).

### Limitations

All assumptions underlying Poiseuille’s law and Ohmic resistances in laminar flows apply. The boundary conditions strongly influence the solution because the system is forced to comply with the preset boundary pressures. However, the effects of a changing diameter on the pressure and flow within the network are evident. We also assume that the empirical formulas to calculate blood viscosity apply to the cerebral microcirculation of mice.

### Statistical analysis

Datasets are presented as mean ± SEM with individual data points. The normality of data was assessed using Shapiro-Wilk and graphical tests. Linear mixed effect (LME) model analysis was employed. Vessel segments (PA, precapillary sphincters, and 1^st^ – 3^rd^ order capillaries) were defined as the fixed effect, whereas the mouse age and particular vasculature were included as random effects as needed. Significant differences were estimated by likelihood ratio tests of the LME model with the fixed effect in question against a model without the fixed effect. Tukey-Kramer’s post hoc test was used for pairwise comparisons between elements in the fixed effect group. All statistical analyses were performed using R studio (version 3.4.4, packages lme4 and multcomp).

