## supplementary text for "Impaired dynamics of brain precapillary sphincters and pericytes at first order capillaries explains reduced neurovascular functions in aging"

**Supplementary Figure 1.** **Comparison of vasodilation elicited by papaverine puff, pinacidil puff, and whisker pad (WP) stimulation.** (**a**) Relative diameter change elicited by the three stimulations in adult and (**b**) old mice. (**c**) Absolute diameter change elicited by the three stimulations in adult and (**d**) old mice. (**e**) Maximal diameter at peak dilation elicited by the three stimulations in adult and (**f**) old mice. Linear mixed effect models were used to test for differences among vessel segments, followed by Tukey post hoc tests for pairwise comparisons. Data are given as mean ± SEM. * indicates p<0.05, ** indicates p<0.001, *** indicates p<0.0001. L1 – L5 denote cortical sublayers.

**Supplementary Figure 2.** **Comparison of whisker pad stimulation-induced vascular responses before and after ET1 puff in adult and aged mice.** (**a**) Comparison of the relative diameter increase by dilation before and after ET1 puff in adult mice and (**b**) old mice. (**c**) Comparison of the relative diameter decrease by undershoot before and after ET1 puff in adult mice and (**d**) old mice. (**e**) Comparison of the maximal diameter at peak dilation before and after ET1 puff in adult mice and (**f**) old mice. Adult: N=5 animals, n=9 vessels. Old: N=7 animals, n=15 vessels. Linear mixed effect models were used to test for differences among vessel segments, followed by Tukey post hoc tests for pairwise comparisons. Data are given as mean ± SEM. * indicates p<0.05, ** indicates p<0.001, *** indicates p<0.0001.

**Supplementary Figure 3. Immunohistochemical analysis of pericyte morphology and α-smooth muscle actin (αSMA) density with aging.** (**a**) Immunohistochemistry of αSMA (green) with endogenesis red indicator in pericytes and smooth muscle cells. The nucleus stained blue (Hoechst). Image processing procedure for the z-stack was to free-hand draw the background area and then calculate the mean background fluorescence intensity for each image plane. The background intensity was normalized to the peak value of the z-stack. Finally, the z-stack images were projected onto one image by maximal projection. (**b**) Two methods were used to measure αSMA intensity. Left: Hand-drawn delineation of the vessel region in the green image and the mean intensity calculated in the selected region. Right: The NG2 image was used to select the pericyte-positive pixels and the mean intensity of the corresponding pixels calculated in the αSMA image. (**c**) Endothelial cell soma density obtained by dividing the number of endothelial cell somas and the examined vessel length. (**d**) NG2 density at the whole vessel surface determined by dividing the NG2 fluorescence intensity and the whole vessel area. (**e**) αSMA density at the whole vessel surface obtained by dividing the αSMA fluorescence intensity and the whole vessel area. Adult: N=3 animals, n=25 vessels. Old: N=3 animals, n=26 vessels. Linear mixed effect models were used to test for differences among vessel segments, followed by Tukey post hoc tests for pairwise comparisons. Data are given as mean ± SEM. * indicates p<0.05, ** indicates p<0.001, *** indicates p<0.0001.

**Supplementary Figure 4.** ***In vivo* vascular structural changes with aging.** (**a** – **c**) Total vessel number per brain volume in adult and old somatosensory cortex. (**a**) General total vessel number, (**b**) capillary order dependent total vessel number, and (**c**) cortical depth dependent total vessel number. (**d** – **f**) Total vessel volume per brain volume in adult and old somatosensory cortex. (**d**) General total vessel volume, (**e**) capillary order dependent total vessel volume, and (**f**) cortical depth dependent total vessel volume. Adult: N=6 animals, n=52 vessels. Old: N=4 animals, n=29 vessels. Linear mixed effect models were used to test for differences among vessel segments, followed by Tukey post hoc tests for pairwise comparisons. Data are given as mean ± SEM. * indicates p<0.05, ** indicates p<0.001, *** indicates p<0.0001. L1 – L5 denote cortical sublayers.
