## Supplementary figures and images for "Impaired dynamics of brain precapillary sphincters and pericytes at first order capillaries explains reduced neurovascular functions in aging"

### supplementary figure 1

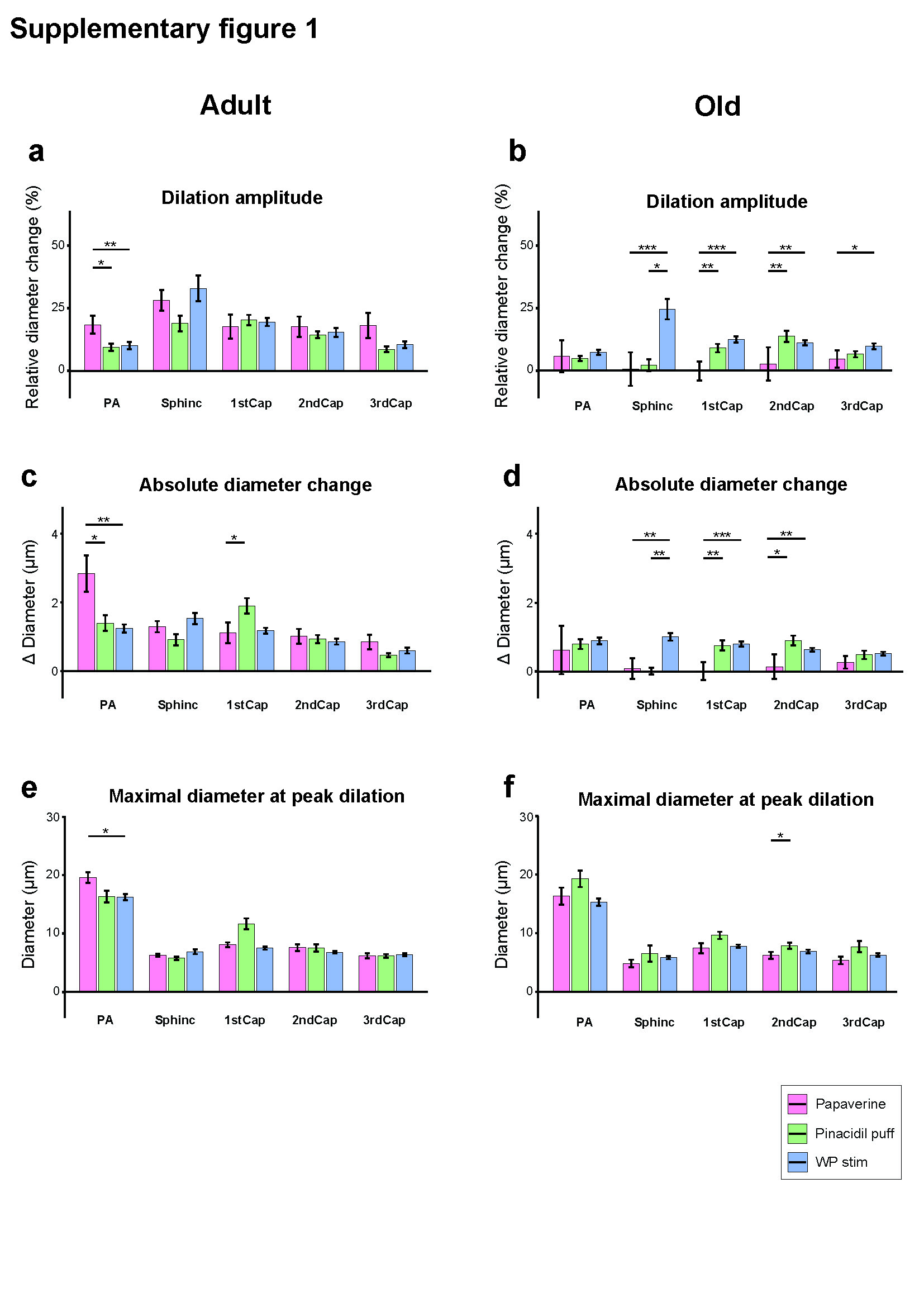

### supplementary figure 2

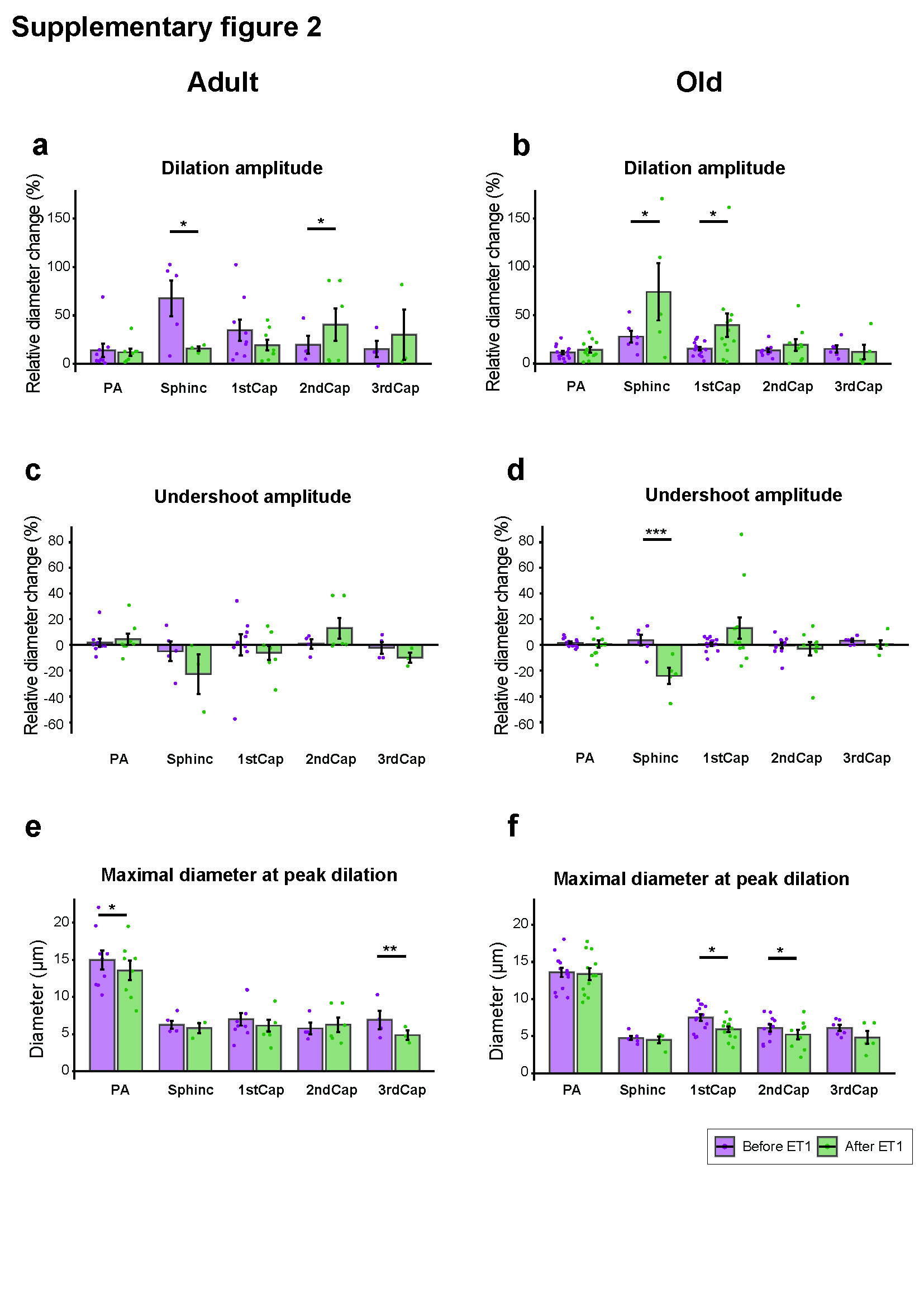

### supplementary figure 3

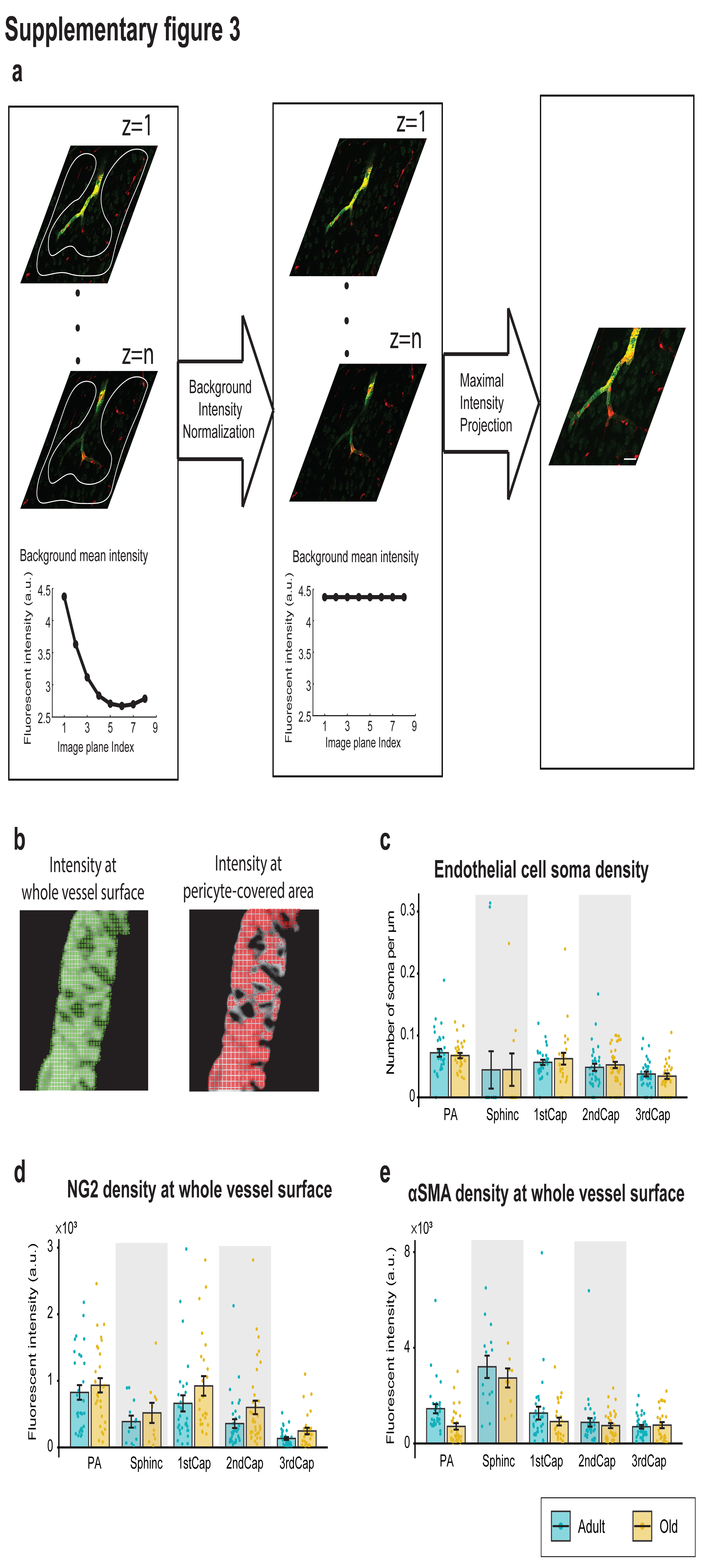

### supplementary figure 4

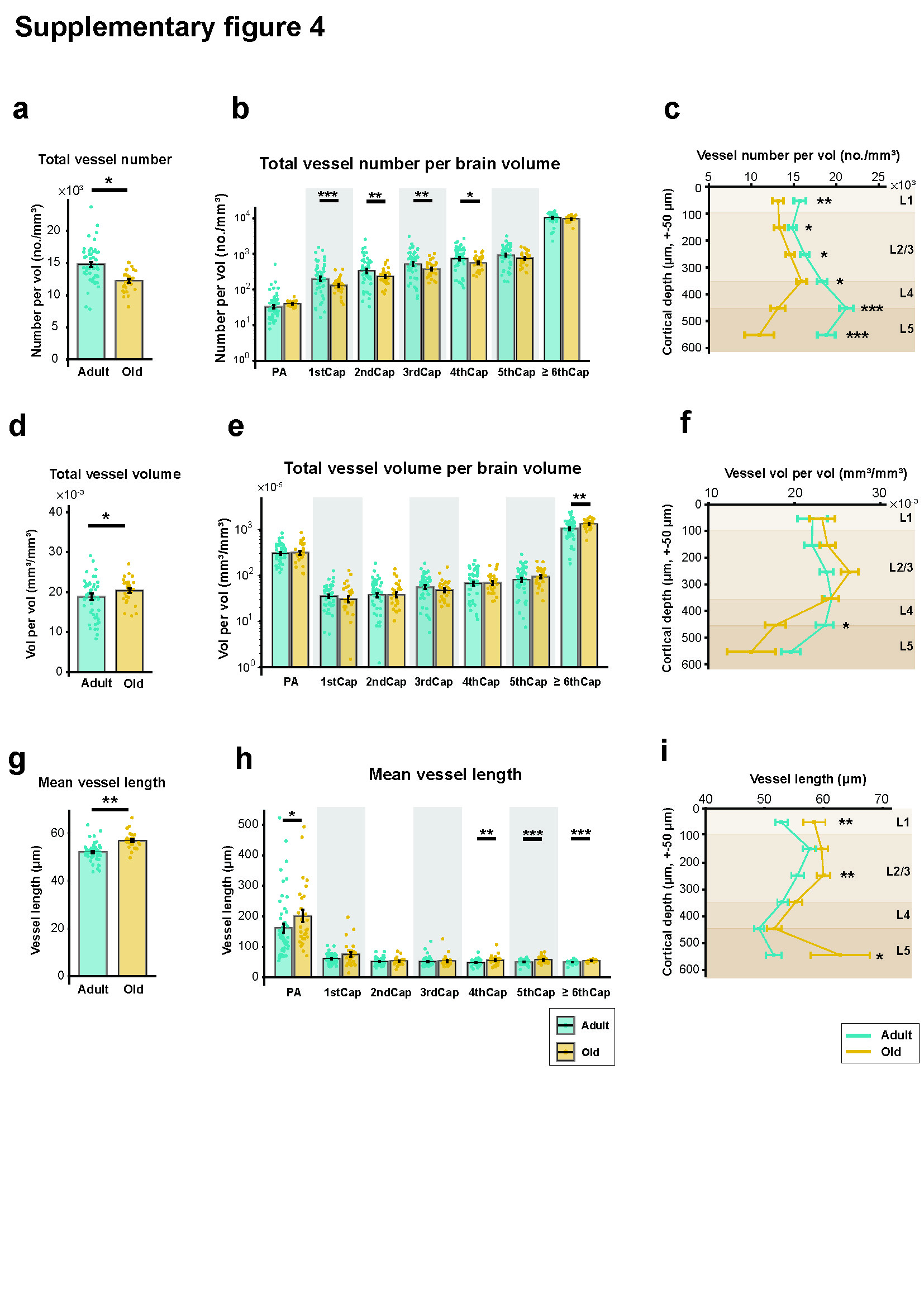
